# Development of a Time-Resolved Fluorescence Resonance Energy Transfer ultra-high throughput screening assay for targeting SYK and FCER1G interaction

**DOI:** 10.1101/2024.06.11.598473

**Authors:** Yuhong Du, Dongxue Wang, Vittorio L. Katis, Elizabeth L. Zoeller, Min Qui, Allan I. Levey, Opher Gileadi, Emory-SAGE-SGC TREAT-AD Center, Haian Fu

## Abstract

The spleen tyrosine kinase (SYK) and high affinity immunoglobulin epsilon receptor subunit gamma (FCER1G) interaction has a major role in the normal innate and adaptive immune responses, but dysregulation of this interaction is implicated in several human diseases, including autoimmune disorders, hematological malignancies, and Alzheimer’s Disease. Development of small molecule chemical probes could aid in studying this pathway both in normal and aberrant contexts. Herein, we describe the miniaturization of a time-resolved fluorescence resonance energy transfer (TR-FRET) assay to measure the interaction between SYK and FCER1G in a 1536-well ultrahigh throughput screening (uHTS) format. The assay utilizes the His-SH2 domains of SYK, which are indirectly labeled with anti-His-terbium to serve as TR-FRET donor and a FITC-conjugated phosphorylated ITAM domain peptide of FCER1G to serve as acceptor. We have optimized the assay into 384-well HTS format and further miniaturized the assay into a 1536-well uHTS format. Robust assay performance has been achieved with a Z’ factor > 0.8 and signal-to-background (S/B) ratio > 15. The utilization of this uHTS TR-FRET assay for compound screening has been validated by a pilot screening of 2,036 FDA-approved and bioactive compounds library. Several primary hits have been identified from the pilot uHTS. One compound, hematoxylin, was confirmed to disrupt the SYK/FECR1G interaction in an orthogonal protein–protein interaction assay. Thus, our optimized and miniaturized uHTS assay could be applied to future scaling up of a screening campaign to identify small molecule inhibitors targeting the SYK and FCER1G interaction.

## 1. Introduction

Spleen tyrosine kinase (SYK) is a major signal transducer in innate and adaptive immune response pathways but is also involved in non-immune biological processes such as cellular differentiation and adhesion[1]. SYK dysregulation occurs in a number of human diseases, including immune thrombocytopenia, autoimmune disorders, and hematological malignancies[2]. More recently, SYK has been implicated in neuroinflammatory conditions, including Alzheimer’s Disease (AD). Interestingly, SYK has demonstrated paradoxical roles in regulating AD pathology. SYK is upregulated in AD individuals, and SYK activity has been positively associated with markers of AD pathology including microglial-mediated neuronal death, Tau protein phosphorylation and accumulation, and Aβ production[3-7]. However, two compelling recent studies in 5xFAD mice, an amyloid plaque heavy model organism, found that loss of SYK activity resulted in increased AD pathology, including higher levels of Aβ and decreased cognitive function in the 5xFAD mouse model[8, 9]. Further study of SYK activity, including the use of chemical probes, is necessary to dissect the role of SYK in AD pathology.

SYK exists naturally in an autoinhibited state with two interdomains (interdomain A and interdomain B) bound to the kinase domain. Key to SYK signaling are protein-protein interactions (PPI) with adaptor proteins with immunoreceptor tyrosine-based activation motifs (ITAMs), in particular B-cell receptors (BCR), Fc receptors (FcR), and C-type lectin receptors (CLR)[1, 2, 10]. ITAMs are short peptide sequences containing at least two tyrosine residues. Phosphorylation of both tyrosine residues permits ITAM binding to the two Src homology 2 (SH2) domains of SYK and thus induces a conformational change freeing the SYK kinase domain. SYK activation can also occur by phosphorylation of the interdomains by Src family kinases, such as Lyn[11-14]. Two hypotheses for modes of activation of SYK have been proposed: the “OR-gate” hypothesis in which either binding of the SH2 domains to a doubly phosphorylated ITAM or phosphorylation of the SYK interdomains activates SYK and the alternate hypothesis in which binding of the phosphorylated ITAM allows a conformation change more permissive to phosphorylation of the interdomains A and B for activation of SYK[10, 13, 15]. Chemical probes could be a key tool in resolving these theories of SYK activation.

In this study, we develop a time-resolved fluorescence resonance energy transfer (TR-FRET) high throughput screening (HTS) assay to monitor the interaction of SYK and high affinity immunoglobulin epsilon receptor subunit gamma (FCER1G), an ITAM-containing subunit common to multiple Fc receptors[16, 17]. FCER1G expression has also been positively associated with AD risk and pathology[18-20]. In this TR-FRET system, we use recombinant protein comprising the tSH2 domain of SYK (residues M6-N269) fused to an N-terminal 6xHis tag (6xHis-SYKtSH2) and a FITC-labeled FCER1G p-ITAM peptide (FITC-p-FCER1G). His-SYKtSH2 is indirectly labeled by terbium (Tb) using an anti-His-Tb antibody. Tb serves as a TR-FRET donor. FITC in FITC-p-FCER1G peptide serves as an acceptor. The interaction of His-SYKtSH2 and FITC-p-FCER1G brings Tb and FITC together and leads to the generation of a TR-FRET signal. We first optimized this TR-FRET assay in a 384-well HTS format and then further minimized the assay into a 1536-well ultra-HTS (uHTS) format. Robust assay performance for screening was achieved in both 384-well HTS and 1536-well uHTS formats. We further conducted a pilot screening of 2,036 compounds and validated the utility of this assay in the 1536-well uHTS format for the discovery of SYK/FCER1G protein-protein interaction (PPI) inhibitors. Screening of large-scale compound libraries using this uHTS assay for SYK and FCER1G interaction could yield potential chemical probes as disruptors of the SYK and FCER1G interaction. These probes could help answer ongoing questions about the mechanism of SYK activity and the role of SYK in human disease.

## 2. MATERIALS AND METHODS

### 2.1 Reagents

The Emory Enriched Bioactive Library (EEBL) of 2,036 FDA approved and bioactive compounds is as previously described[21]. Anti-His terbium conjugated antibody is a mouse monoclonal antibody (Anti 6His-Tb cryptate, Cat. No. 61HISTLF, Cisbio Bioassays). Anti-Flag d2 conjugated antibody was purchased from Cisbio Bioassays (Cat. 61FG2DLF). Hematoxylin was purchased from MedChem Express (Cat. No. HY-N0116, Batch 19245, CAS No. 517-28-2). Expression and purification of recombinant proteins and peptides were as described previously[22].

### 2.2 Cell culture and conditions

HEK293T (ATCC, CRL-3216) cells were cultured in Dulbecco’s modified Eagle’s medium (DMEM, Corning, 10-013-CV) with 10% fetal bovine serum (Sigma, F0926-500ML), 100 units penicillin and 100 μg streptomycin (Sigma, P0781-100ML) at 5% CO_2_ and 37°C in a humid environment.

### 2.3 Cloning and constructs

GST-tagged full-length SYK and VenusFlag-tagged full-length FCER1G were generated using Gateway LR Clonase (Invitrogen, 11791-100) for mammalian cell expression. Vector pDONR223-SYK for GST-tagged full-length SYK was purchased from Addgene (Plasmid # 23907). Vector pDONR221-FCER1G for vFlag-tagged full length FCER1G was purchased from DNASU (Clone ID HsCD00075915). Plasmids, including maps and sequences, created in this manuscript are available through Addgene: pDEST-GST-hSYK (Plasmid #211892) and pcDNA3-hFCER1G-VenusFlag (Plasmid #211899).

### 2.4 Assay development in 384-well HTS and 1536-well uHTS format

The SYKtSH2 and FCER1G-ITAM TR-FRET assay was carried out in 384-well black solid bottom plates (Corning Costar, 3573) with total volume of 25 μL per well and miniaturized in 1536-well black solid bottom plates (Corning Costar, 3724) with a total volume of 5 μL. All experiments were conducted in FRET buffer (20mM Tris, pH 7.0, 0.01% Nonidet P40, and 50mM NaCl). FRET buffer purified His-SYKtSH2 protein, FITC-FCER1G peptide, and anti-His Tb (1:1000) were added to plates simultaneously. Plates were incubated at room temperature, sealed from light in aluminum foil for indicated times. TR-FRET signal was then read by the PHERAstar FSX plate reader (BMG LABTECH). TR-FRET signal is calculated by ratio of the emission fluorescence of the acceptor fluorophore (FITC) divided by the donor fluorescence (F520nm/F490nm) and multiplying by 10,000.

### 2.5 Assay performance evaluation for HTS

Z’ factor was calculated to determine the quality of the assay for HTS in the absence of compound [23]. For the positive and negative controls, 16 wells in the 384-well HTS format and 32 wells in the 1536-well uHTS format were analyzed. Z’ = (3*SD_b_ + 3*SD_f_) / (F_b_ – F_f_), where F_b_ is mean TR-FRET signal from His-SYKtSH2, anti-His-Tb, and FITC-p-FCER1G alone (bound peptide) and SD_b_ is the standard deviation of this signal. F_f_ is the mean TR-FRET signal from anti-His Tb and FITC-p-FCER1G alone (free peptide in background conditions, and SD_f_ is the standard deviation of this signal. A Z’ factor between 0.5 and 1 indicates a robust assay that is suitable for HTS or uHTS. Signal to background ratio, S/B, was calculated as F_b_/F_f_. S/B is an indicator of assay sensitivity and the dynamic window of the assay.

### 2.6 uHTS pilot screen

To validate the assay for screening in a 1536-well uHTS format, a pilot screening was carried out using the EEBL library containing 2,036 FDA approved and bioactive compounds. In brief, the reaction mixture as optimized was dispensed at 5 μL per well into 1536-well black plates. Then 0.1 μL of library compound (1 mM) dissolved in DMSO was added using pintool integrated with Beckman NX (Beckman Coulter, Brea, CA). The final compound concentration was 20 μM and the final DMSO concentration was 2%. After incubating at 4 °C for 16 h, TR-FRET signals were measured using PHERAstar FSX plate reader as already described. Screening data were analyzed using Bioassay software from CambridgeSoft (Cambridge, MA). The S/B and Z’ in uHTS format were calculated. The effect of compound on the interaction TR-FRET signal was expressed as percentage of control and calculated as the following equation: % of control = ((FRETsample – FRETblank) / (FRETcontrol – FRETblank))*100%, where FRETsample is the FRET signal from sample condition, FRETblank is the FRET signal from negative control wells not containing active FRET components, and FRETcontrol is FRET signal from His-SYKtSH2, anti-His Tb, and FITC-p-FCER1G alone.

### 2.7 GST pull-down and cell-lysate full length TR-FRET assay

HEK293T cells were transfected with plasmid containing FCER1G (full length)-VenusFlag (Addgene Plasmid #211899) and GST-SYK (full-length) (Addgene Plasmid #211892) in a 6-well plate (Costar® 6-well Clear TC-treated Multiple Well Plates, 3516, Corning) at 1*10^6 cells per well. 1.5 μg FCER1G-VenusFlag, 1.5 μg GST-SYK, and 9μg PEI were incubated with 2 mL DMEM medium at room temperature for 30 min before being added drop. 48 hours after transfection, cells were harvested by pipetting on ice and transferred to pre-cold Eppendorf tube. Cells were then pelleted for 3 min at 2000 rcf and washed once with PBS. Cell pellets were finally lysed (5 sec sonication in ice bath) with SYK lysis buffer (25 mM HEPES pH7.5, 200 mM NaCl, 0.5% Triton-X 100, and protease inhibitors, Pierce Protease Inhibitor Mini Tablets EDTA free, Thermo Fisher, A32955), Phosphatase Inhibitor Cocktail 2 (Sigma, P5726), and Phosphatase Inhibitor Cocktail 3 (Sigma, P0044)).

For GST pull-down, 180 μL diluted cell lysates (1:3 dilution) with 20 μL Glutathione Sepharose 4B beads added (17075605, Cytiva) were incubated with inhibitors or DMSO only controls for 2 hours on a rotator at 4°C. Beads were washed by lysis buffer for 3 times and boiled in 2X Laemmli Sample buffer (Bio-RAD, #1610737). Monoclonal ANTI-FLAG M2-HRP antibody (Sigma A8592) at 1:1000 and primary antibody GST-Taq (2.6H1) Mouse mAb, (Cell Signaling 2624S) at 1:1000 with secondary antibody peroxidase-conjugated AffiniPure Goat Anti-Mouse IgG (H+L) (Jackson Immuno Research,115-035-003) at 1:5000 were used for immunoblotting.

For the full-length PPI TR-FRET assay development, cells were lysed as above after over-expressing GST-SYK and FCER1G (full length)-VenusFlag by transfection methods described above. Cell lysates were diluted 40X in FRET buffer (20mM Tris, pH 7.0, 0.01% Nonidet P40, and 50mM NaCl). 5 μL of diluted lysates were added into a 1536-well black polystyrene plate (Corning 3724). Diluted antibodies, anti-GST-Tb (1:1000, Fisher Scientific, 50-211-7733) and anti-Flag-d2 (1:500 dilution, Cisbio Bioassays Cat. 61FG2DLF), were added at 0.2 μL per well by robotic dispenser. The plate was incubated at 4°C overnight. The TR-FRET signal for full length PPI was detected with the PHERAstar FSX plate reader as described above.

### 2.8 Bio-Layer Interferometry (BLI)

Bio-Layer Interferometry (BLI) was performed using the Octet RED384 system (Sartorius BioAnalytical Instruments). The biotinylated SYK-tSH2 (50 μg/mL, C-terminal AviTag, N-terminal his tag cleaved) was loaded onto Octet Super Streptavidin Sensors (Sartorius 18-5057) in 50 μL 1XBLI kinetic buffer (Octet Kinetics Buffer 10 X, 10 X KB, Sartorious PN:18-11-5, diluted in 1XPBS) for 10 minutes resulting in a 14 nm loading density. After loading, the Octet Super Streptavidin Sensors were associated with compound in gradient concentration in assay buffer (20 mM HEPES pH7.5, 200 mM NaCl, 0.005% Tween20, 1 mM TCEP and 4% DMSO). Then the sensors/compound were dissociated in assay buffer. Dissociation constant (K_d_) = ([SYK-tSH2] [Compound]) / [Compound, SYK-tSH2 complex]. K_d_ and K_d_ error were analyzed by ForteBio Data Analysis 9.0.

## 3. RESULTS

### 3.1 Design of TR-FRET assay for SYK and FCER1G interaction

Fluorescence resonance energy transfer (FRET) can occur between two fluorophores with overlapping emission and excitation spectra. When the donor fluorophore is excited, the donor emits an energy within the excitation spectrum of the acceptor fluorophore, which, if in near proximity to the donor, then emits a signal readable in an assay format. Using a donor fluorophore with a long-lived fluorescence allows for temporal delay between excitation of the donor and detection of the acceptor emission (TR-FRET). This time-delayed feature minimizes the background interference of fluorescence from short-lived sources, such as medium and compound fluorescence.

To develop the TR-FRET assay for SYK and FCER1G interaction, we used recombinant protein comprising of the tSH2 domain of SYK (residues M6-N269) fused to an N-terminal 6xHis tag (6xHis-SYKtSH2) and a FITC-labeled FCER1G p-ITAM peptide (FITC-p-FCER1G). The 6xHis tag of the SYKtSH2 is bound to terbium (Tb) using Tb labeled anti-His antibody (anti-His-Tb). When His-SYKtSH2 binds to the FITC-p-FCER1G, the terbium is brought close to the FITC. Excitation of the terbium (337 nm laser) leads to emission of Tb (490 nm) within the excitation spectrum of FITC, thereby generating a TR-FRET signal measured at 520 nm (Fig. 1).

**Figure 1.**
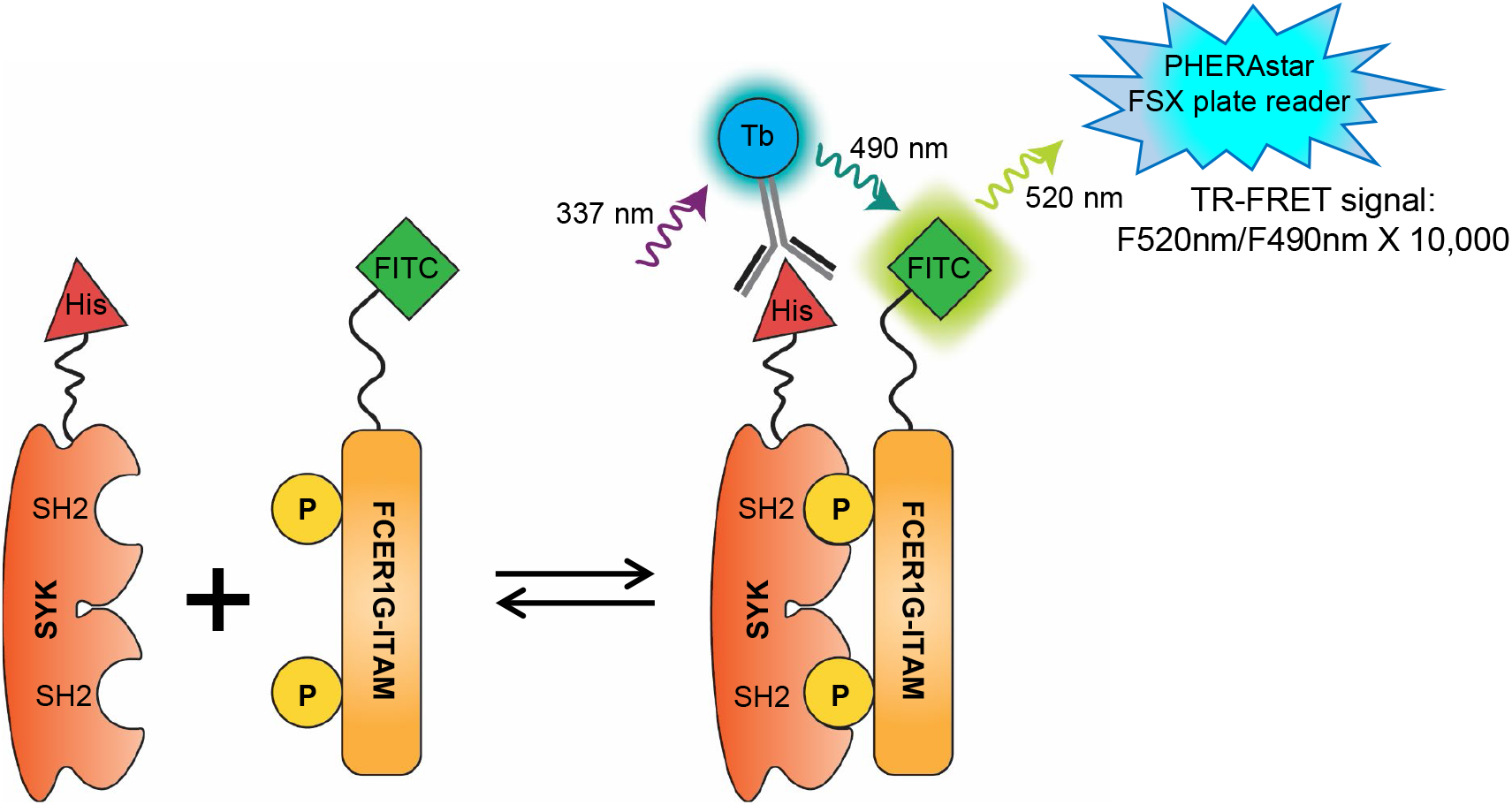
Design of TR-FRET assay for SYK and FCER1G interaction. The SH2-domains of SYK are labeled with a His protein antigen (His-SYKtSH2) which is detected by a terbium (Tb) labeled anti-His antibody. The phosphorylated ITAM domain of FCER1G is labeled with fluorescein isothiocyanate (FITC-p). When the SH2 domains of SYK and the phosphorylated sites of the FCER1G ITAM domain interact, the fluorophores are brought into close proximity. Upon excitation at 337 nm, donor fluorophore Tb emits energy at 490 nm that transfers to acceptor FITC, which emits a 520 nm FRET signal to be read by the plate reader.

### 3.2 Development of SYK/FCER1G TR-FRET in 384-well HTS format

To optimize SYKtSH2 and FCER1G-ITAM interaction TR-FRET assay, we first carried out two ways titration by mixing increasing concentrations of His-SYKtSH2 protein with increasing concentrations of p-FCER1G peptide in a 384-well HTS format. As shown in Fig. 2A, incubating increasing concentrations of FITC-p-FCER1G peptide with various doses of His-SYHtSH2 led to dose-dependent increases of signal to background ratio (S/B) of TR-FRET signal. Similarly, titration of His-SYHtSH2 with various doses of FITC-p-FCER1G peptide led to dose-dependent increases of S/B of TR-FRET signal (Fig. 2B). Based on the protein and peptide titration results, the HTS reaction mixture containing 2 nM His-SYKtSH2, 6 nM FITC-p-FCER1G, and anti-His-Tb (1:1000 dilution), which generates about EC_80_ of the maximal TR-FRET signal was selected for the further evaluation. As shown in Fig. 2C, the HTS reaction mixture generated significant TR-FRET signal compared to background control with FITC-p-FCER1G peptide and anti-His-Tb only (Fig. 2C). The TR-FRET signal is robust in 384-well HTS format with Z’ factor greater than 0.8 (Fig. 2D) and the signal to background ratio (S/B) greater than 12 (Fig. 2E). The signal is stable from 10 minutes to overnight (Fig. 2E).

**Figure 2.**
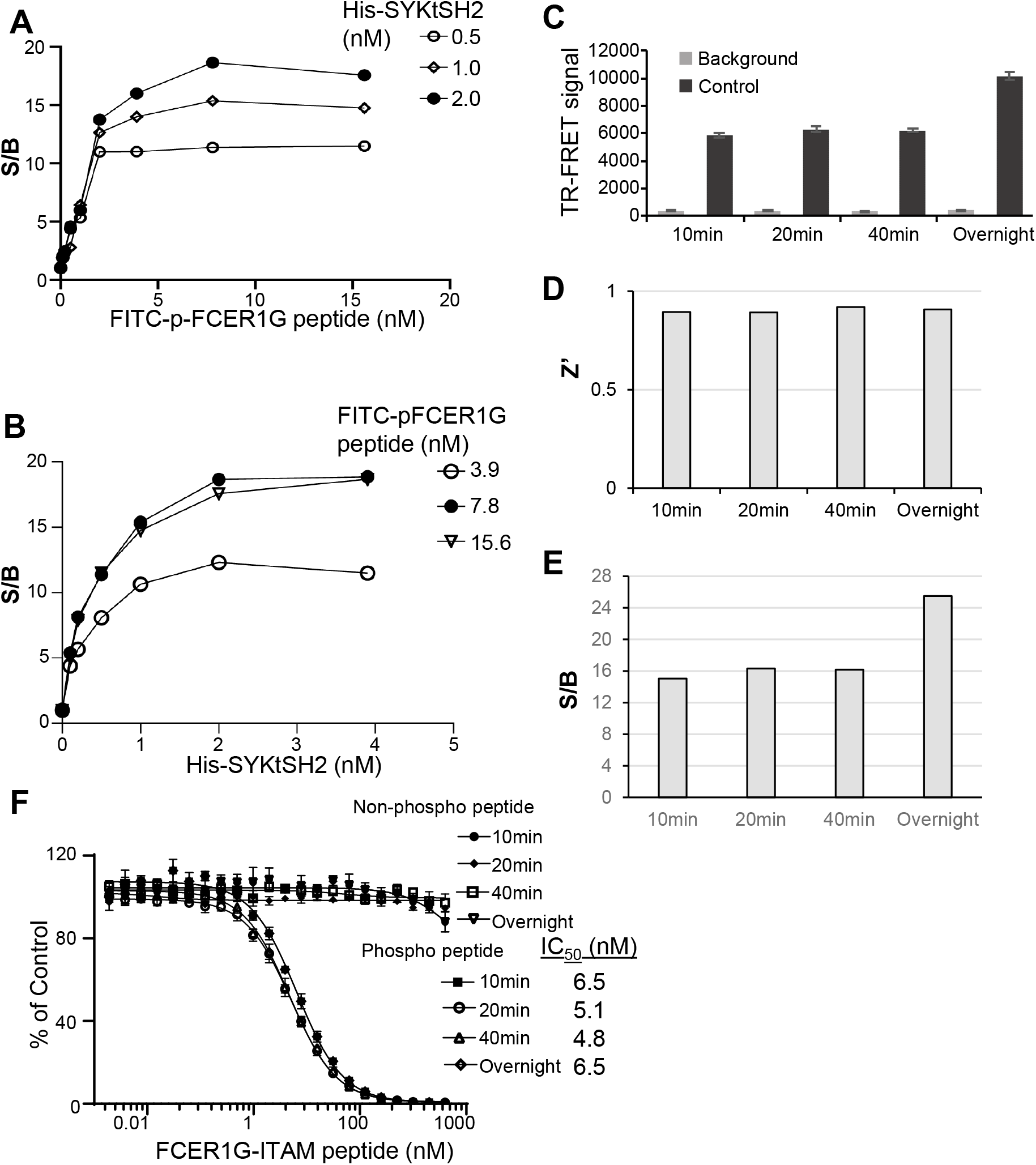
Assay performance of SYK/FCER1G TR-FRET in 384-well HTS format. A) Titration of FITC-p-FCER1G peptide with His-SYKtSH2. B) Titration of His-SYKtSH2 with FITC-p-FCER1G peptide. C) TR-FRET signal for background and control conditions across multiple timepoints. Background conditions omit His-SYKtSH2. Data shown are averages with SD from 16 replicates for each sample. D) Z’ factor is greater than 0.8 across all timepoints. E) Signal to background ratio (S/B) is greater than 12 across all timepoints. F) Comparison of activity of unlabeled, non-phosphorylated and unlabeled, phosphorylated FCER1G-ITAM peptides in competition with labeled, phosphorylated FCER1G-ITAM by TR-FRET across timepoints. Error bars represent SD from triplicate samples.

To validate the TR-FRET assay for screening inhibitors of SYK/FCER1G, we tested the inhibitory effect of unlabeled FCER1G peptide on His-SYKtSH2/FITC-p-FCER1G TR-FRET signal. The reversibility of labeled peptide binding is important when testing potential disruptors of the PPI as the assay must be sensitive to decreases in signal. For this experiment, the HTS reaction mixture (2 nM His-SYKtSH2, 1:1000 anti-His Tb, and 6 nM FITC-p-FCER1G) served as the control. The background conditions included only 6 nM FITC-p-FCER1G and 1:1000 anti-His Tb. The bound labeled FITC-p-FCER1G peptide was out competed for binding with His-SYKtSH2 with increasing concentrations of the unlabeled FCER1G-phospho-peptide. The TR-FRET signal decreased in a dose-dependent manner as concentrations of the unlabeled FCER1G-phospho-peptide increased (Fig. 2F). The unlabeled FCER1G-phospho-peptide demonstrated an IC_50_ of 5 nM. This inhibitory effect on TR-FRET signal was also stable over 10 minutes, 20 minutes, 40 minutes, and overnight. An unlabeled, unphosphorylated FCER1G peptide was used as a negative control and did not have an effect on the TR-FRET signal produced by the His-SYKtSH2 and FITC-p-FCER1G interaction. Thus, this 384-well HTS assay is specifically sensitive to disruption and is suitable for screening SYK/FCER1G PPI inhibitors.

### 3.3 Miniaturization of SYK/FCER1G TR-FRET into 1536-well uHTS format

In efforts to increase throughput and reduce reagent costs, we further miniaturized the TR-FRET assay into a 1536-well uHTS format. A matrix containing 15-point titrations of His-SYKtSH2 and FITC-p-FCER1G peptide were performed using 5 μL/well in a 1536-well format. TR-FRET signal patterns were very similar to those obtained with the 384-well format. We selected the same conditions of HTS reaction buffer used with 384-well format (2 nM His-SYKtSH2, 1:1000 anti-His Tb, and 6 nM FITC-p-FCER1G) for further assay validation. As shown in Fig. 3A, the reaction mixture in 1536-well format was scaled down to 5 μL per well (Fig. 3A). The uHTS also performed well with increases in TR-FRET signal for control over background at 40 minutes and 18 hours (Fig. 3B). The 1536-well uHTS assay generated robust signal with a Z’ factor greater than 0.7 and a S/B greater than 15 (Fig. 3C,D), indicating a high-quality assay for uHTS. Similar with results in the 384-well plate format, the none-labeled FCER1G-phospho-peptide was able to compete the TR-FRET signal in a dose-dependent manner with IC_50_ about 5 nM in 1536-well uHTS format, and the effect was stable from 40 minutes to overnight. These data demonstrate that the uHTS assay is robust and sensitive for compound screening.

**Figure 3.**
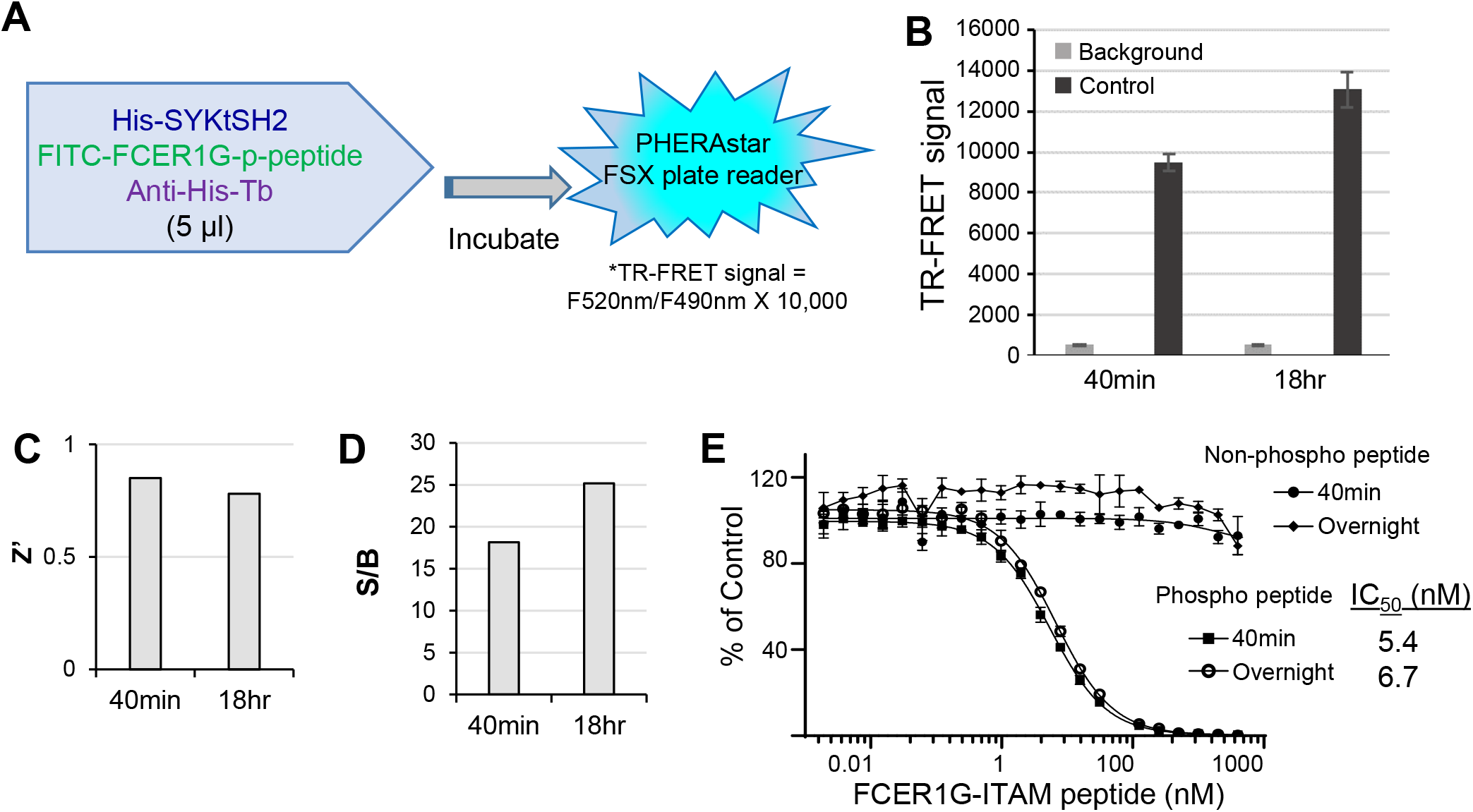
Assay performance of SYK/FCER1G TR-FRET in 1536-well uHTS format. A) Outline of experimental procedure for 1536-well format. His-SYKtSH2 (2 nM), anti-His Tb (1:1000), and phosphorylated FITC-FCER1G (6 nM) are added at 5 μL final volume per well before incubation and subsequent reading of TR-FRET signal. B) TR-FRET signal for background and control conditions at 40 min and 18 hr. Data shown are averages with SD from 32 replicates for each sample. Background conditions omit His-SYKtSH2. C) Z’ factor is greater than 0.7 across both timepoints. D) Signal to background ratio (S/B) is greater than 15 across both timepoints. E) Comparison of activity of unlabeled, non-phosphorylated and unlabeled, phosphorylated FCER1G-ITAM peptides in competition with labeled, phosphorylated FCER1G-ITAM by TR-FRET across timepoints. Data are averaged with SD across 4 replicates per sample.

### 3.4 Pilot screening in 1536-well uHTS format

To validate the uHTS assay performance, we performed a pilot screen with 2,036 FDA approved and bioactive compounds library at 20 μM final across six 1536-well plates. The TR-FRET signal across the six plates was consistent (Fig. 4A). Robust signal has been achieved with Z’ greater than 0.6 and S/B greater than 10 for all six plates (Fig. 4B,C). Screening results are shown in Fig. 4D. 26 compounds have been identified as primary hits with % of Control < 70 as hit cut-off. The hit rate is 1.3%.

**Figure 4.**
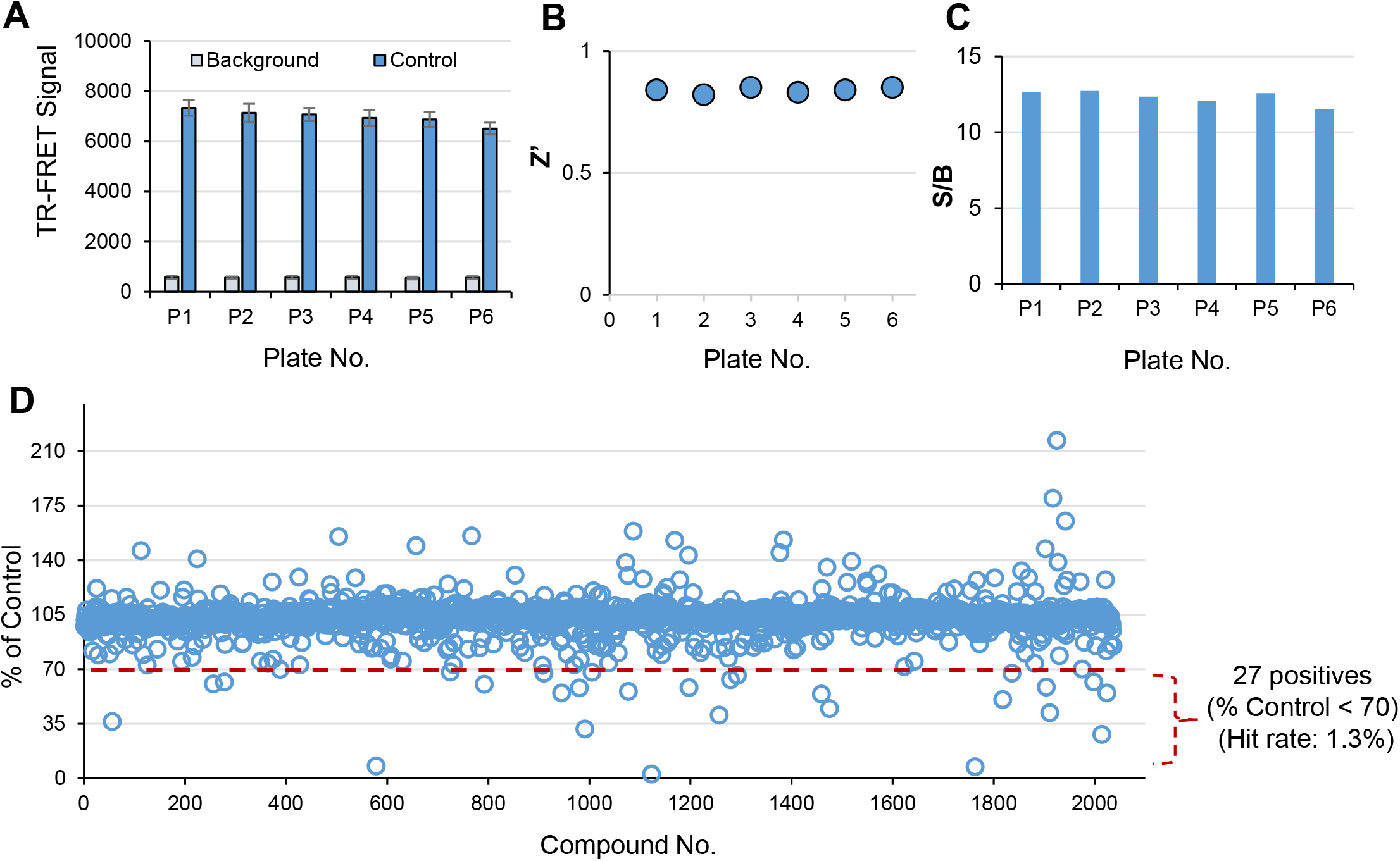
Pilot screening in 1536-well uHTS format for SYK/FCER1G inhibitors. A) TR-FRET signal is consistent across all six 1536-well plates, error bars represent SD for n=32 replicates for each sample. B) Z’ factor is greater than 0.6 across all plates. C) S/B is greater than 10 across all plates. D) Scatter plot of pilot screening of 2,036 FDA approved and bioactive compounds in 1536-well uHTS format.

### 3.5 Validation of hit compounds

The 26 positive hits from pilot uHTS were subjected to a dose-response (DR) TR-FRET assay. 13 compounds demonstrated an IC_50_ less than 30 μM from library stock compounds. These 13 compounds were reordered and confirmed in dose-response TR-FRET assay. One compound, hematoxylin (CAS No. 517-28-2) (Fig. 5A), decreased the TR-FRET signal in a dose-dependent manner with IC_50_ at 1 μM (Fig. 5B). Hematoxylin was further evaluated in a GST pull-down assay for the ability to disrupt the interaction between full length SYK and FCER1G in a cell lysate. Cell lysates from HEK 293T cells co-expressing full-length GST-tagged SYK and full-length Flag-tagged FCER1G were mixed with increasing concentrations of compound. The GST-tag of SYK was pulled down by immunoprecipitation. The Flag-tag of FCER1G was then detected by Western blot to determine the association of FCER1G with SYK in the presence of compound. Hematoxylin disrupted the SYK/FCER1G interaction in a dose-dependent manner by the GST pulldown (Fig. 5C). Dose-response TR-FRET was also conducted with full length SYK and FCER1G constructs and yielded an IC_50_ of 9 μM (Fig. 5D). Direct binding between hematoxylin and SYK protein were then measured. Bio-Layer Interferometry (BLI) utilizing immobilized biotinylated SYKtSH2 protein demonstrated direct binding of hematoxylin to SYK with dissociation constants (K_d_) of 28.1 μM (Fig. 5E). These data have validated hematoxylin as a promising SYK/FCER1G PPI inhibitor.

**Figure 5.**
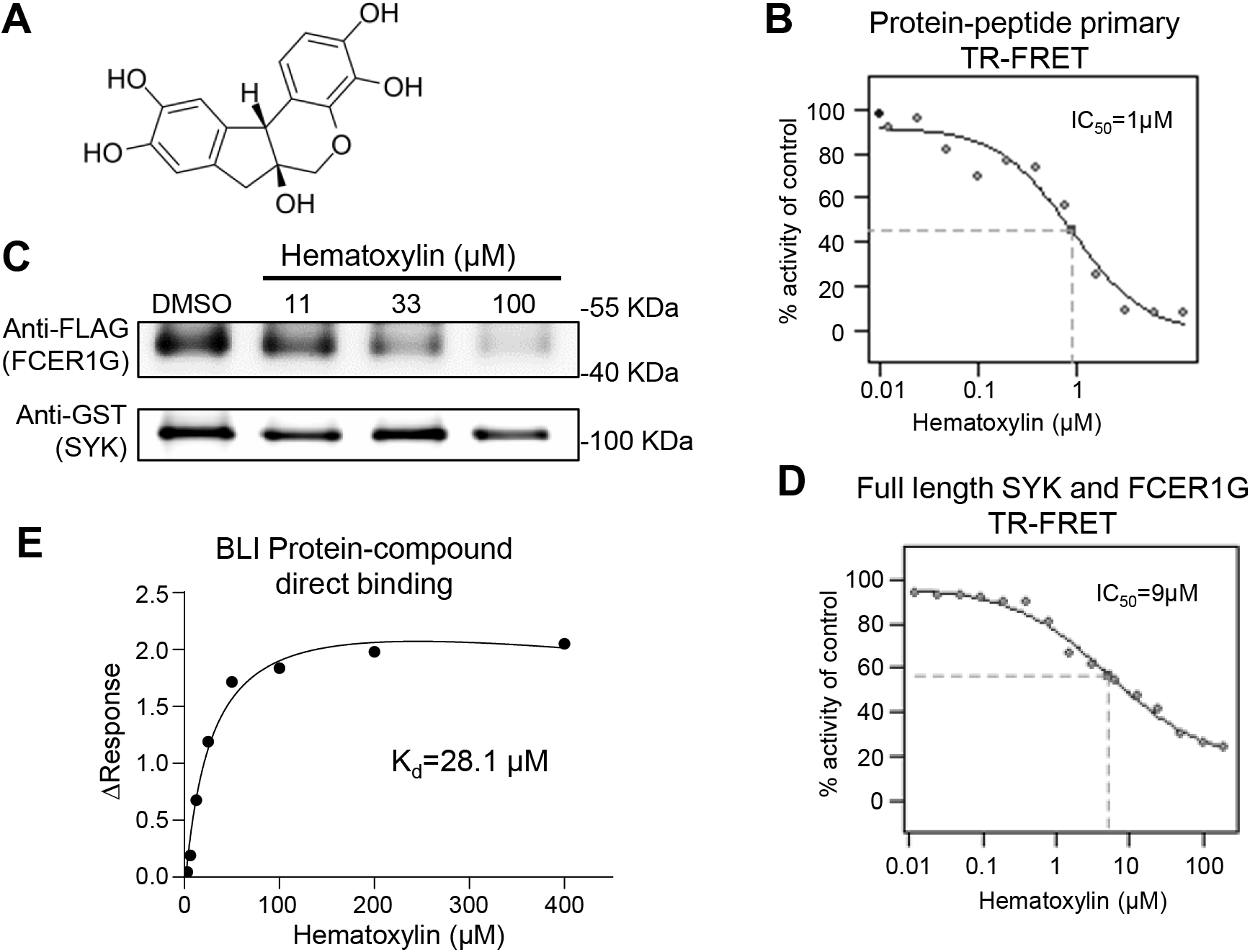
Validation of hematoxylin as SYK/FCER1G PPI inhibitors. A) Structure of hematoxylin. B) Dose-dependent inhibition of hematoxylin on primary TR-FRET assay. Data shown averaged from 4 replicates for each sample. C) Hematoxylin exhibited inhibitory effect in GST-pull down assay with full length SYK and FCER1G PPI. Cell lysates expressing both constructs were incubated with indicated concentration of compound before pulling down the GST tag (SYK) and detecting associated vFlag (FCER1G) by Western blot. D) Hematoxylin dose-dependently inhibited the TR-FRET signal of full length SYK and FCER1G PPI using cell lysate. Data shown averaged from 4 replicates for each sample. E) Bio-Layer Interferometry (BLI) for hematoxylin interaction with biotinylated SYKtSH2 protein direct binding with K_d_=28.1 μM.

## 4. Discussion

SYK has emerged as a potential target for several human diseases, including Alzheimer’s Disease. Currently, one SYK inhibitor, fostamatinib, has been approved by the FDA for treatment of immune thrombocytopenia, an autoimmune disease affecting platelets and clotting[24]. Fostamatinib and the other SYK inhibitors in clinical trials rely almost exclusively on inhibiting kinase activity as a mechanism of action[2, 25]. However, SYK activity can also be regulated through its capacity for autoinhibition. Binding of phosphorylated ITAM-containing proteins to the SH2 domains of SYK induces a conformational change to promote activity of the SYK kinase domain. Additionally, phosphorylation of SYK by the Src family kinases activates SYK. Binding of phosphorylated ITAM by SYK and Src-mediated phosphorylation of SYK could occur sequentially or as separate SYK activating models. Use of a chemical probe altering the SYK ITAM interaction could help resolve these models of SYK activation and increase our understanding of how SYK activity is regulated[13]. Herein, we report the development and validation of a high-throughput screen assay to identify modulators of the SYK/phospho-FCER1G-ITAM interaction.

To monitor the SYK/FCER1G PPI, we utilized a time-resolved fluorescence resonance energy transfer (TR-FRET) technique as previously described to increase specificity of signal by minimizing background signal[22]. We first developed the assay in a 384-well HTS format with a final volume of 25 μL per well. This 384-well format assay demonstrated suitable parameters for HTS with a Z’ greater than 0.8 and S/B values greater than 12. We further miniaturized the assay into a 1536-well uHTS format with a final volume of 5 μL per well. The 1536-well uHTS has similarly robust performance with a Z’ greater than 0.7 and S/B values greater than 15.

To validate the utilization of the uHTS assay for compound screening, we performed a pilot uHTS of 2,036 FDA approved and bioactive compounds library in 1536-well format. A panel of secondary assays has confirmed and validated one top compound, hematoxylin, as a promising inhibitor for SYK/FCER1G PPI. Hematoxylin exhibited an IC_50_ of 1 μM in primary DR TR-FRET assay. It showed disruption of full length SYK/FCER1G interaction in both GST-pulldown and TR-FRET with protein over-expressed cell lysate. BLI compound/protein direct binding assay has shown that it directly binds to SYK protein. Interesting, hematoxylin, a natural organic compound most often associated as a dye for cytology and histology, has been reported to inhibit amyloid β-protein fibrillation with an IC_50_ of 1.6 μM, alleviate amyloid-Induced cytotoxicity, and inhibit tau protein aggregation[26, 27].

In summary, we have developed and validated a TR-FRET assay for screening small molecule inhibitors for the SYK/FCER1G interaction. We have used this validated TR-FRET assay for expanded large scale screening of 138,214 diversity compounds library for SYK/FCER1G PPI inhibitors[22]. Furthermore, similar configurations of our uHTS assay can be applied to determine the specificity of PPI inhibitors for phosphorylated ITAM domains of binding partners with SYK in addition to FCER1G, leading to the potential for context or cell-type specific SYK inhibitors[10]. Of particular interest would be studies of how hits identified could alter phosphorylation of SYK by Src kinases and kinase activity of SYK. Overall, the development and validation of TR-FRET uHTS assay will enable and allow for efficient screening campaigns to identify SYK inhibitors for biology and therapeutic development.

## Abbreviations

6xHis-SYKtSH2: tSH2 domain of SYK (residues M6-N269) fused to an N-terminal 6xHis tag
AD: Alzheimer’s Disease
anti-His-Tb: Tb labeled anti-His antibody
BCR: B-cell receptors
BLI: Bio-Layer Interferometry
CLR: C-type lectin receptors
DR: dose-response
EEBL: Emory Enriched Bioactive Library
FCER1G: immunoglobulin epsilon receptor subunit gamma
FcR: Fc receptors
FITC-p-FCER1G: FITC-labeled FCER1G p-ITAM peptide
HTS: high throughput screening
ITAM: immunoreceptor tyrosine-based activation motif
K^d^: dissociation constant
PPI: protein-protein interaction
S/B: signal-to-background
SH2: Src homology 2 domain
SYK: spleen tyrosine kinase
Tb: terbium
TR-FRET: time-resolved fluorescence resonance energy transfer
uHTS: ultra-high throughput screening

## Contributions

Y.D., D.W., V.L.K., M.Q., A.I.L, O. G, and H.F. conceptualization; Y.D., D.W., V.L.K., M.Q., A.I.L, O. G, and H.F. methodology; Y.D., D.W., and M.Q. formal analysis; Y.D., D.W., and M.Q. investigation; V.L.K. and O.G. resources; E.L.Z. writing-original draft; Y.D., D.W., V.L.K., M.Q., E.L.Z., A.I.L, O. G, and H.F. writing-review and editing; Y.D., D.W., M.Q., and E.L.Z. visualization; A.I.L., H.F., and Y.D. supervision; E.L.Z. project administration.

## Acknowledgements

The research reported in this manuscript was led by the Emory-Sage-SGC TREAT-AD center and supported by the National Institute on Aging grant U54AG065187. The Target Enablement to Accelerate Therapy Development for Alzheimer’s Disease (TREAT-AD) Consortium was established by the National Institute on Aging (NIA). Members of Emory-Sage-SGC TREAT-AD Center include: Ishita Ajith^1^, Joel K. Annor-Gyamfi^2^, Jeff Aube^2^, Alison D. Axtman^2^, Frances M. Bashore^2^, Ranjita S. Betarbet^3^, Juan Botas^4^, William J. Bradshaw^1^, Paul E. Brennan^1^, Peter J. Brown^2^, Robert R. Butler 3rd^5^, Jacob L. Capener^2^, Gregory W. Carter^6^, Gregory A. Cary^6^, Catherine Chen^4^, Rachel Commander^3^, Sabrina Daglish^2^, Suzanne Doolen^7^, Yuhong Du^3^, Duc M. Duong^3^, Aled M. Edwards^8,^*, Michelle E. Etoundi^4^, Kevin J. Frankowski^2^, Stephen V. Frye^2^, Haian Fu^3^, Opher Gileadi^1^, Marta Glavatshikh^2^, Jake Gockley^9^, Katerina Gospodinova^1^, Anna K. Greenwood^9^, Peter A. Greer^10^, Lea T. Grinberg^11^, Shiva Guduru^2^, Levon Halabelian^8^, Crystal Han^5^, Brian Hardy^2^, Laura M. Heath^9^, Stephanie Howell^2^, Hasi Huhe^7^, Andrey A. Ivanov^3^, Suman Jayadev^12^, Vittorio L. Katis^1^, Stephen Keegan^6^, May Khanna^13^, Dmitri Kireev^2^, Carl LaFlamme^14^, Karina Leal^9^, Hyemin Lee^3^, Tom V. Lee^4^, Tina M. Leisner^2^, Allan I. Levey^3,*^, Qianjin Li^3^, David Li-Kroeger^4^, Zhandong Liu^4^, Benjamin A. Logsdon^9^, Frank M. Longo^5^, Lara M. Mangravite^9^, Peter S. McPherson^14^, Richard M. Nwakamma^3^, Felix O. Nwogbo^2^, Carolyn A. Paisie^6^, Arti Parihar^12^, Kenneth H. Pearce^2^, Kun Qian^3^, Min Qui^3^, Stacey J Sukoff Rizzo^7^, Karolina A. Rygiel^1^, Julie Schumacher^5^, David D. Scott^15^, Nicholas T. Seyfried^3^, Joshua M. Shulman^4^, Benjamin Siciliano^3^, Arunima Sikdar^2^, Nathaniel Smith^4^, Michael Stashko^2^, Judith A. Tello Vega^15^, Dilipkumar Uredi^2^, Dongxue Wang^3^, Jianjun Wang^3^, Xiaodong Wang^2^, Zhexing Wen^3^, Jesse C. Wiley^9^, Alexander Wilkes^1^, Charles A. Williams^12^,Timothy M. Willson^2^, Aliza Wingo^3^, Thomas S. Wingo^3^, Novak Yang^3^, Jessica E. Young^12^, Miao Yu^6^, Elizabeth L. Zoeller^.3^ *Leads: Aled M. Edwards and Allan I. Levey, contact, respectively.

^1^University of Oxford, Oxford, OX3 7FZ, UK

^2^University of North Carolina at Chapel Hill, Chapel Hill, NC 27599, USA

^3^Emory University School of Medicine, Atlanta, GA 30322, USA

^4^Baylor College of Medicine, Houston, TX 77030, USA

^5^Stanford University School of Medicine, Stanford, CA, 94305, USA

^6^The Jackson Laboratory, Bar Harbor, ME 04609, USA

^7^University of Pittsburgh School of Medicine, Pittsburgh, PA 15219, USA

^8^University of Toronto, Toronto, ON M5G 1L7, Canada

^9^Sage Bionetworks, Seattle, WA, 98121, USA

^10^Queen’s University, Kingston, Ontario, ON K7L 3N6, Canada

^11^University of California, San Francisco, San Francisco, CA 94143, USA

^12^University of Washington, Seattle, WA 98109, USA

^13^New York University, New York, NY 10010, NY, USA ^14^McGill University, Montreal, QC H3A 2B4, Canada ^15^University of Arizona, Tucson, AZ 85724, USA

